# Structure-function coupling reveals hemisphere-specific network reorganization after stroke

**DOI:** 10.64898/2026.09.18.752042

**Authors:** F. Mahani, G.R. Fink, M. Hoehn, M. Aswendt

**Author notes:** **Correspondence to:** Prof. Dr. Markus Aswendt, Department of Neurology, University Hospital Frankfurt, Theodor-Stern-Kai 7, D-60590 Frankfurt am Main. equal senior authors (M. Hoehn, M. Aswendt).

## Abstract

MRI is an excellent tool for monitoring functional and structural network reorganization after brain injury and may provide biomarkers for treatment stratification and novel neuromodulatory interventions. In ischemic stroke, functional network changes, particularly within the contralesional hemisphere, and structural integrity, most prominently of the corticospinal tract, have been successfully associated with behavioral deficit and outcome. However, these components are commonly studied separately and at single time points, which does not reflect the highly dynamic nature of brain network plasticity. Here, we characterized structural and functional connectivity (SC-FC) coupling in the healthy mouse brain and longitudinally assessed intra- and inter-hemispheric connections up to 4 weeks after cortical and cortico-striatal stroke. In the healthy brain, intra-hemispheric SC-FC coupling was lateralized in the sensorimotor network (SMN), whereas the default mode network (DMN) showed no significant hemispheric difference. Intra-hemispheric coupling exceeded inter-hemispheric coupling in the SMN. After cortical stroke, regions with high baseline intra-hemispheric SC-FC coupling remained largely stable, whereas initially weakly coupled sensorimotor regions showed delayed increases within the ischemic hemisphere. Contralesional intra-hemispheric SC-FC coupling changed little, but directional inter-hemispheric analyses revealed selective remodeling, including increased contralesional-to-ischemic coupling of primary motor cortex and reduced bidirectional coupling of medial sensorimotor cortex. Cortico-striatal stroke produced fewer longitudinal changes, dominated by reduced inter-hemispheric motor-cortex coupling. These alterations did not simply correspond to regional lesion involvement. Thus, hemisphere- and direction-resolved SC-FC coupling uncovers network- and stroke-model-specific reorganization not detectable when structural and functional connectivity are analyzed independently.

**Significance statement:** Recovery after stroke is believed to depend on sufficient functional and structural brain network reorganization over time, yet these processes are usually studied separately. We show that their relationship, quantified as structure–function coupling, follows distinct organizational principles in the healthy mouse brain and is selectively remodeled after stroke. Coupling differed between sensorimotor and default mode networks and between intra- and interhemispheric connections. After stroke, changes depended on region, hemisphere, connection direction, and lesion model and did not simply correspond to local tissue damage. These findings identify hemisphere- and direction-resolved structure–function coupling as a complementary measure of poststroke network reorganization and provide a framework for testing its relationship with recovery and network-targeted interventions.

## INTRODUCTION

Brain function emerges from interactions within structurally constrained neuronal networks, yet structural and functional connectivity provide complementary rather than interchangeable descriptions of network organization. Diffusion-weighted MRI (DWI) and resting-state functional MRI (rs-fMRI) enable these two components to be studied noninvasively in humans and, increasingly, in experimental animal models (1). The widely accepted notion of a close relationship between both methods was based on reasonable evidence from healthy brain analysis that “function follows form”, i.e., that functional connections follow the underlying structural neuronal networks (2). Experimental studies in rodents further supported this concept. In mice, rs-fMRI functional connectivity (FC) between homotopic cortical and hippocampal regions and along cortico-striatal pathways largely corresponded to monosynaptic tracer connectivity, whereas FC within some subcortical circuits emerged despite the absence of direct anatomical connections (3). However, these studies examined healthy brains, and how regional structure-function relationships reorganize longitudinally after brain injury remains largely unknown

A new concept to quantitatively describe the relationship between structural connectivity and FC is structure-function (SC-FC) coupling, calculated as the correlation between the strength of both network systems (4–6). SC-FC coupling varies across networks and brain states and altered values have been reported in neurological diseases (6). Studies in stroke patients provided the first evidence that SC-FC coupling in capsular compared to pontine stroke is weaker, despite FC and SC disruption in both (7), and that lower SC-FC coupling is related to larger motor deficits (8). This pioneering work with stroke patients highlighted that focal lesions induce widespread network disruption, including SC-FC coupling changes in the healthy, contralesional hemisphere, and that regional or sub-network measures are more informative than a single global value (7–10). Related investigations in animal models exploring SC-FC coupling dynamics by stroke type through repeated imaging across the recovery period are largely lacking (11).

Here, we focused on changes in SC-FC coupling using longitudinal diffusion tensor imaging (DTI) and resting-state fMRI in two mouse stroke models: photothrombosis (PT), which produces a local cortical stroke, and middle cerebral artery filament occlusion (MCAO), which causes a larger cortico-striatal stroke lesion (12, 13). We re-analyzed these datasets to establish pre-stroke SC-FC coupling in the healthy, adult mouse brain within the sensorimotor (SMN) and default mode networks (DMN). Furthermore, we examined how SC-FC coupling changes at the subnetwork and regional levels when analyzed separately for the ipsilesional and contralesional hemispheres, and whether distinct coupling patterns differentiate cortical from cortico-striatal stroke. This approach revealed distinct intra- and interhemispheric organizational features of SC-FC coupling and identified contralateral and interhemispheric remodeling as a major source of stroke-model-specific network differences.

## MATERIALS AND METHODS

Details on the experimental procedures are contained in the *SI Appendix*.

### Study design and animal model

The datasets and experimental procedures were based on previously reported stroke imaging studies and were re-analyzed in a harmonized way to enable direct comparison between diffusion MRI-derived structural measures and resting-state fMRI-derived functional measures (12, 13). All procedures complied with the German Animal Welfare Act and followed the ARRIVE and IMPROVE recommendations (14, 15). Approval was granted by the Landesamt für Natur, Umwelt und Verbraucherschutz North Rhine-Westphalia (approval numbers: 84-02.04.2014.A305, issued on 2015-01-26, and 84-02.04.2016.A461, issued on 2017-07-06, for work performed at the Max Planck Institute for Metabolism Research, as well as 81-02.04.2019.A309, issued on 2019-11-25, for work performed at the University Hospital Cologne. Animals were maintained on a 12 h light/dark schedule with unrestricted access to food and water. Cortical stroke was induced by photothrombosis in N = 22 adult male C57BL/6J mice (8-12 weeks old, body weight 25 - 30 g, The Jackson Laboratory). Cortico-striatal stroke was induced by 30 min middle cerebral artery occlusion (MCAO) in N = 6 adult male NMRI-nu mice (aged 12-14 weeks, 32-37 g, Janvier Labs). In both cohorts, T2-weighted MRI, diffusion-weighted MRI, and resting-state fMRI were performed before stroke induction (baseline) and repeated 1, 2, and 4 weeks later. This longitudinal design allowed us to evaluate SC and FC over the same recovery period within a unified atlas-based framework.

### Quantification of SC-FC coupling

We calculated SC-FC coupling separately for each subject at each time point, using a row-wise correlation approach similar to that described previously (4). From the structural and functional connectivity matrices, we extracted 4 corresponding matrix compartments: intra-hemispheric connections within the ischemic hemisphere, intra-hemispheric connections within the contralateral hemisphere, inter-hemispheric connections from ischemic to contralateral, and inter-hemispheric connections from contralateral to ischemic. Within each matrix compartment, we determined SC-FC coupling for each brain region by calculating the Pearson correlation coefficient between the corresponding rows of the SC and FC matrices. This row-wise correlation quantified how closely the structural connectivity profile of a given region matched its functional connectivity profile, yielding one coupling value per region for each hemisphere-specific or inter-hemispheric compartment (Fig. 2A). To assess longitudinal changes, the resulting coupling values were averaged across subjects at each time point, which provided a time-resolved measure of SC-FC coupling for each region and each hemispheric connection type. The analyses focused on regions belonging to the sensorimotor network and the default mode network, both of which are relevant to stroke-related network reorganization (16, 17).

**Figure 1:**
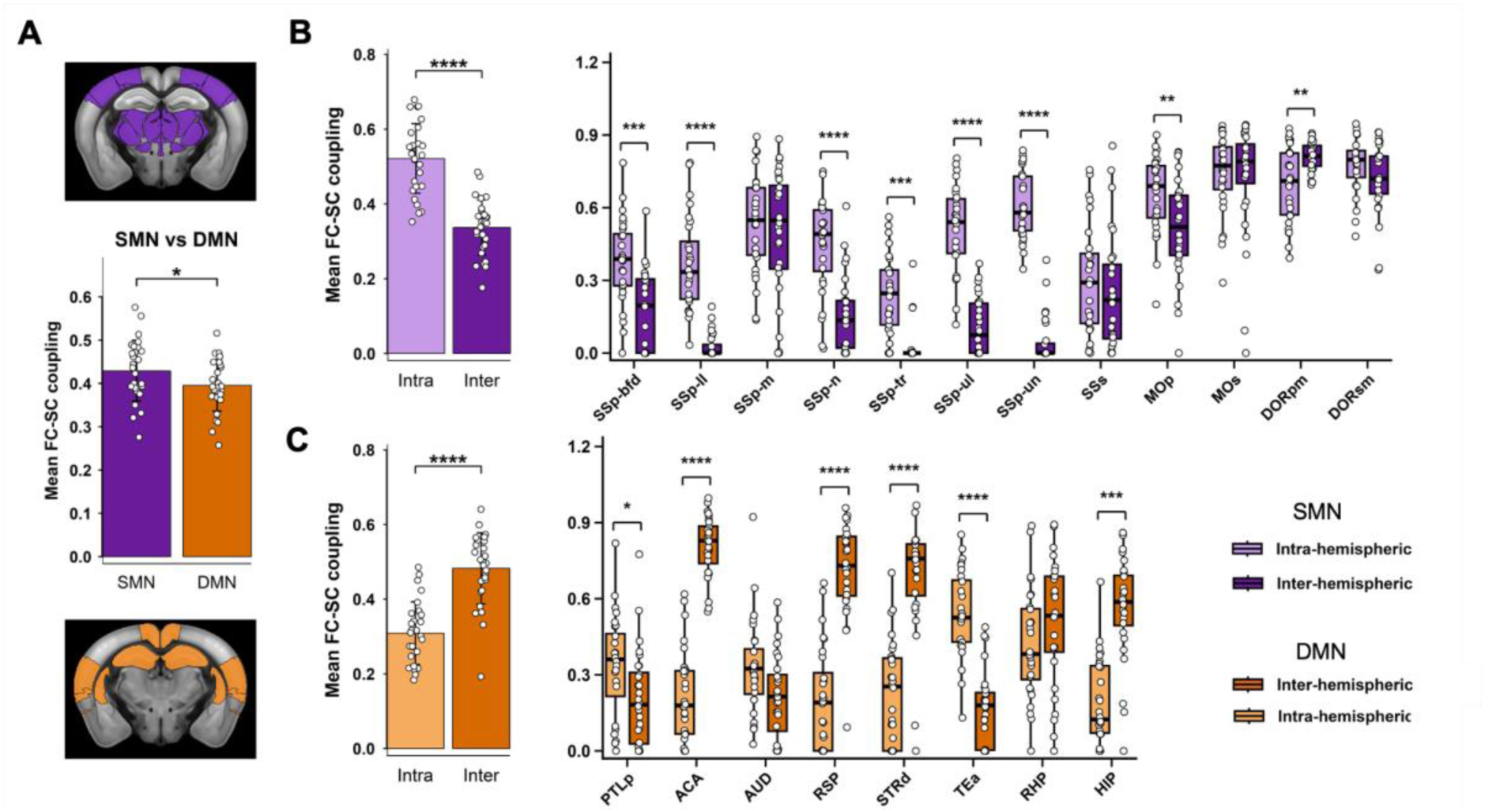
Intra- and inter-hemispheric structure–function coupling (SC-FC coupling) within the sensorimotor (SMN) and default mode (DMN) networks. **A)** SC-FC coupling comparison between SMN (purple) and DMN (orange) sub-networks (visualized as atlas regions - top and bottom, respectively), showing higher SC-FC coupling in SMN (p = 0.020, paired-samples t-test). **B)** SMN: Left, mean intra-hemispheric SC-FC coupling was significantly higher compared to inter-hemispheric SC-FC coupling averaged across regions (p < 0.0001, paired-samples t-test with Benjamini–Hochberg false discovery rate correction). Right, analyses of individual SMN components comparing intra-hemispheric (light purple) and inter-hemispheric (dark purple) coupling were performed using paired Wilcoxon signed-rank tests with Benjamini–Hochberg false discovery rate correction across regions. Intra-hemispheric coupling was significantly higher in SSp-bfd, SSp-ll, SSp-n, SSp-tr, SSp-ul, SSp-un, and MOp, whereas inter-hemispheric coupling was significantly higher in DORpm. **C)** DMN: Left, mean intra-hemispheric and inter-hemispheric SC-FC coupling across the entire DMN. Whole-network intra-hemispheric and inter-hemispheric couplings were compared using a paired-samples t-test with Benjamini–Hochberg false discovery rate correction. Right, we performed regional comparisons using paired Wilcoxon signed-rank tests with Benjamini– Hochberg false discovery rate correction across regions. The whole-network significance symbols are based on FDR-adjusted paired t-tests, whereas the regional significance symbols are based on FDR-adjusted paired Wilcoxon signed-rank tests. Bars represent mean ± SD, box plots show the median, interquartile range, and full data distribution, and circles indicate individual animals (both stroke groups pooled). Statistical significance is indicated as p < 0.05 (*), p < 0.01 (**), p < 0.001 (***), and p < 0.0001 (****).

**Figure 2:**
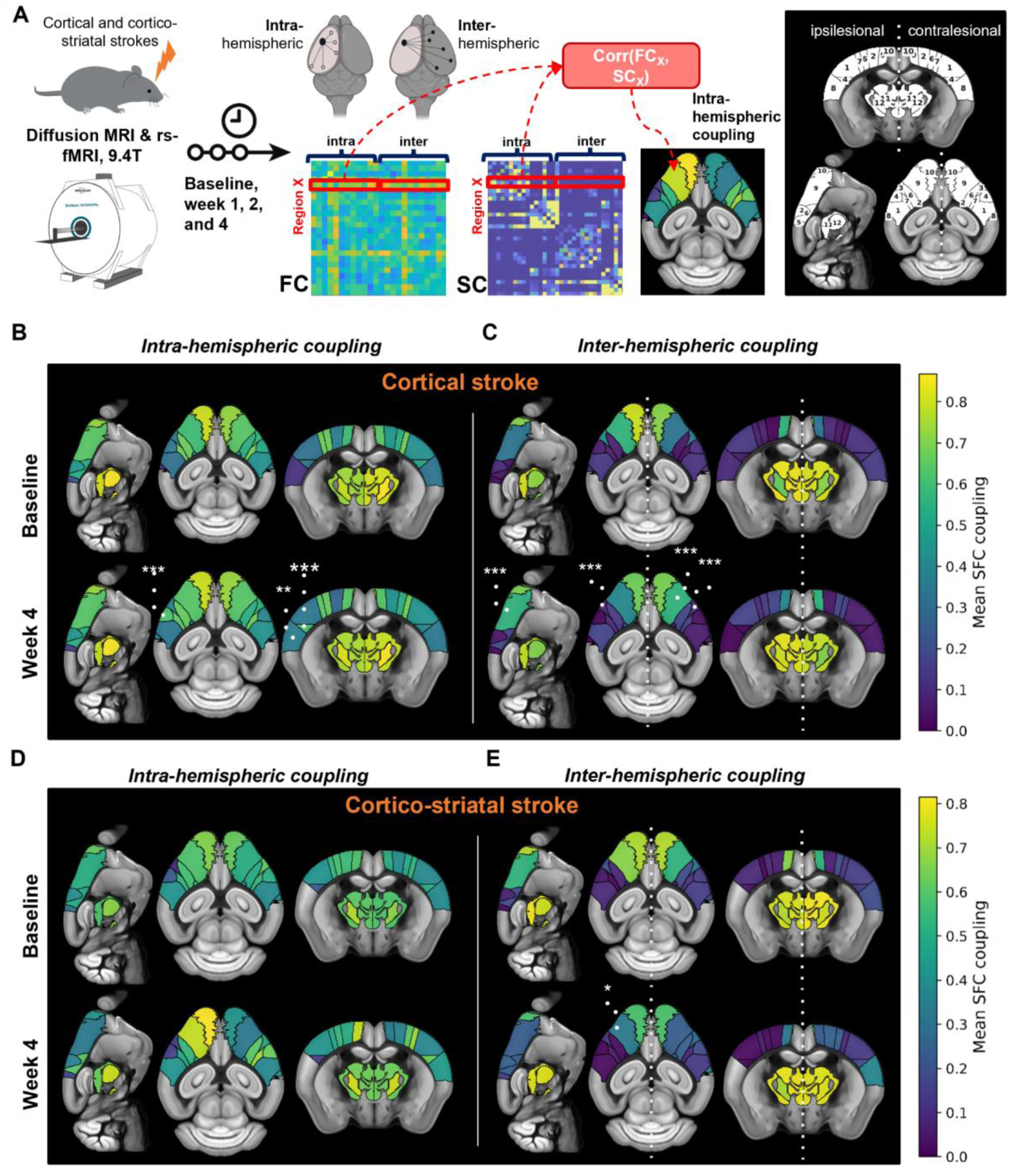
SC-FC coupling changes in the sensorimotor network 4 weeks after stroke. **A)** Experimental paradigm and strategy to calculate SC-FC coupling from region-wise FC and SC data separately for the left and right hemisphere. Right, illustration of selected brain regions in the SMN: 1) SSp-bfd, 2) SSp-ll, 3) SSp-m, 4) SSp-n, 5) SSp-tr, 6) SSp-ul, 7) SSp-un, 8) SSs, 9) MOp, 10) MOs, 11) DORpm, 12) DORsm. **B-D)** SC-FC coupling changes in SMN regions overlaid on the Allen Mouse Brain template; mean SC-FC coupling colored (0-purple to 0.8 yellow). SC-FC coupling maps are shown for intra- and inter-hemispheric coupling in cortical and cortico-striatal stroke. For each stroke model and coupling direction, longitudinal SC-FC coupling changes were analyzed using linear mixed-effects models with Day, Region, and their interaction as fixed effects and MouseID as a random intercept. Estimated marginal means were used to compare each post-stroke time point with baseline within each region, followed by Benjamini–Hochberg false discovery rate correction across all Day-versus-baseline regional comparisons. Significance symbols shown on the week 4 maps indicate FDR-adjusted week 4-versus-baseline comparisons. Statistical significance is indicated as p < 0.05 (*), p < 0.01 (**), and p < 0.001 (***). SC-FC coupling.

### Statistical analysis

Longitudinal changes in SC-FC coupling relative to baseline were analyzed using a linear mixed-effects model in R (v4.4.1) with *dplyr* for data handling and preprocessing as well as the *lmer* function from the *lme4* package. The model was specified as: Coupling ∼ Day × Region + (1 | MouseID); where Coupling was the dependent variable, Day and Region were included as fixed effects, and MouseID was included as a random intercept to account for repeated measurements within the same animal. We used the interaction term to test whether temporal changes in coupling differed across regions. We performed post hoc pairwise comparisons between time points within each region using estimated marginal means from the *emmeans* package, with Tukey adjustment to control for multiple comparisons.

## RESULTS

### Healthy brain: intra- and inter-hemispheric coupling differences

For baseline (pre-stroke) comparison, we first assessed whether SC-FC coupling differed between mice later allocated to the cortical and cortico-striatal stroke groups. We detected no significant group differences at the combined or network level (***SI Appendix,* Fig. S1A-B**). Likewise, no significant group differences remained after correction at the region- and connection-specific level (***SI Appendix,* Fig. S1C**). Because baseline group differences were absent, we pooled data from both cohorts for subsequent baseline analyses. We then assessed hemispheric differences in baseline SC-FC coupling separately for intra-hemispheric coupling, inter-hemispheric coupling, and a composite measure (***SI Appendix,* Fig. S2**). In the default mode network (DMN), we established equivalence for the composite and directional inter-hemispheric measures. For intra-hemispheric SC-FC coupling, the left-right difference was small (mean difference left - right = −0.023, 90% CI [−0.059, 0.013]), but equivalence within the predefined ±0.05 bounds was not formally established (TOST p = 0.105), with no evidence of a left–right difference in the conventional paired test (p = 0.295). In contrast, in the sensorimotor subnetwork (SMN), the composite and directional inter-hemispheric measures were also equivalent. Intra-hemispheric SC-FC coupling showed a pronounced hemispheric asymmetry and was significantly higher in the right than in the left hemisphere (mean difference = −0.078, 90% CI [−0.112, −0.045]; TOST p = 0.919; paired t-test p < 0.001).

The direct comparison of DMN and SMN revealed significantly higher SC-FC coupling in the SMN (0.43 ± 0.07 vs. 0.40 ± 0.06; p = 0.020; **Fig. 1A**). The average intra-hemispheric SC-FC coupling value for SMN was significantly higher compared to the inter-hemispheric values (avg. 0.52 ± 0.18 vs. 0.33 ± 0.30; p < 0.0001; **Fig. 1B**). Region-wise statistics revealed that the higher intra-hemispheric SC-FC coupling was mainly driven by sensorimotor regions (SSp-ll, SSp-n, SSp-tr, SSp-ul, SSp-un; **Fig. 1C**), which showed the largest difference in intra- vs. inter-hemispheric SC-FC coupling (all: p < 0.0001). Overall, intra-hemispheric SC-FC coupling across regions varied between the SSp-tr 0.242 ± 0.162 (SSp-tr) and 0.778 ± 0.116 (DORsm), with lower values in sensorimotor regions (SSp-tr/ll/n, SSs) and significantly higher values in the thalamus and motor cortex (***SI Appendix,* Fig. S3**). inter-hemispheric SC-FC coupling range was 0.027 ± 0.083 (SSp-tr) to 0.811 ± 0.059 (DORpm), again with lower values in sensorimotor regions (all except for SSp-m) and significantly higher values in the thalamus and motor cortex (***SI Appendix*, Fig. S3**).

With the exception of SSp-m, SSs, MOs, and DORpm/sm, the intra-hemispheric SC-FC coupling values were substantially higher compared to their inter-hemispheric counterparts (**Fig. 1B**). Only in DORpm, intra-hemispheric SC-FC coupling was significantly lower compared to inter-hemispheric SC-FC coupling (0.698 ± 0.155 vs. 0.811 ± 0.059; p = 0.0013). In the DMN, this organization was reversed: mean interhemispheric coupling exceeded intrahemispheric coupling (0.48 ± 0.25 vs. 0.31 ± 0.12; P < 0.0001; Fig. 1C), with particularly low intra-hemispheric values in ACA, RSP, STRd, HIP (0.196 - 0.330), and with higher coupling values in PTLp, TEa, RHP (0.334 - 0.542). Inter-hemispheric couplings were significantly increased in all regions except AUD and RHP, with the highest values ranging from 0.560 to 0.807 (ACA, RSP, STRd, HIP). Only in PTLp and TEa was this reversed (p = 0.033 and p < 0.0001).

Taken together, these findings demonstrate sub-network-specific SC-FC coupling patterns in the healthy mouse brain. SC-FC coupling was higher in SMN than in DMN, and the two networks showed opposite intra- and inter-hemispheric coupling profiles.

### Dynamic SC-FC-coupling changes after stroke

SC-FC coupling was quantified longitudinally at baseline and weeks 1, 2, and 4 post-stroke for the SMN and DMN regions, and separately for the two stroke groups (Fig. 2, matrices in ***SI Appendix*, Fig. S5**).

After cortical stroke, the group of regions (SSp-m, MOp, MOs, DORpm/sm) with comparably high intra-hemispheric SC-FC coupling at baseline remained unchanged (**Fig. 2B, *SI Appendix*, Fig. S5A**). In contrast, SSp-n and SSs, two regions with low SC-FC coupling at baseline, increased first at 2 weeks (p < 0.001) and also at 4 weeks post-stroke (p = 0.002 and p = 0.007, respectively). This difference was not related to the initial extent of stroke (e.g., DORpm/sm: 0%; SSp-n: 76.005±34.243; SSs: 0.515±1.574). No significant longitudinal changes were detected within the contralateral hemisphere (**Fig. 2B, *SI Appendix*, Fig. S5C**). However, directional inter-hemispheric analyses revealed a significant increase in contralateral-to-ischemic MOp coupling at 2 and 4 weeks post-stroke (p = 0.027 and p = 0.005). In contrast, contralateral-to-ischemic SSp-m coupling decreased significantly at 4 weeks (p = 0.005). This reduction in SSp-m coupling was also evident in the ischemic-to-contralateral direction at 2 and 4 weeks (p = 0.005 and p < 0.0001, respectively; **Fig. 2C; *SI Appendix,* Fig. S5G**). In other regions, which also started with very low inter-hemispheric SC-FC coupling, (i.e., most sensorimotor regions except SSp-m), bidirectional coupling increased slightly but not significantly (**Fig. 2C; *SI Appendix*, Fig. S5E,G**). Other regions with initially high inter-hemispheric SC-FC coupling (MOs, DORpm/sm) tended to decrease over time, especially for contralateral-to-ischemic connections. Only ischemic-to-contralateral MOs coupling also decreased significantly at week 1 (p=0.0398).

In the cortico-striatal group, we detected less significant SC-FC coupling changes over time (**Fig. 2D-E, *SI Appendix*, Fig. S5B,D**). Most regions showed only minor fluctuations over time (especially DORpm/sm, SSp-bfd, SSp-ll, SSp-ul, SSp-un). The strongest changes related to baseline, however, did not reach statistical significance; for example, SSp-tr, MOp, and MOs (ipsilateral increase vs. contralateral decrease), and SSp-n, SSp-un, and DORsm (ipsilateral decrease and contralateral increase). Inter-hemispherically, significant SC-FC coupling changes over time were detected in the ischemic to contralateral MOp at week 1 (decrease, p = 0.012) and 4 (decrease, p = 0.022), but not in contralateral to ischemic MOp connections at week 4 (decrease, p = 0.170). Additional trends were observed in ischemic to contralateral MOs (decrease at week 4) and in contralateral to ischemic SSp-ll/ul and SSs (increase at week 4). As in cortical strokes, SC-FC coupling changes were not related to the extent of stroke-induced damage in these regions (***SI Appendix*, Fig. S6**), e.g., SSp-tr (0.622±0.716) and SSp-un (60.290±43.636).

In the DMN, there were only subtle SC-FC coupling changes (***SI Appendix*, Fig. S4**). PTLp decreased contralaterally after cortical stroke and ipsilateral AUD increased at week 4 after cortico-striatal stroke (p = 0.0189 and p = 0.0228). Inter-hemispherically, STRd, the only region among ACA, RSP, and HIP with high SC-FC coupling at baseline, decreased significantly at week 4 post cortical stroke - contralateral to ischemic (p = 0.0013). In cortico-striatal stroke, contralateral to ischemic SC-FC coupling for RHP and HIP transiently decreased at week 2 (p = 0.024 and p = 0.0071). The observed coupling changes did not simply correspond to regional lesion involvement (***SI Appendix*, Fig. S6**), as these regions were only partially affected, such as RHP (10.112 ± 15.141), or showed no detectable lesion involvement, including AUD, RSP, and HIP. In contrast, SC-FC coupling remained stable in regions with substantial lesion involvement, including ACA after cortical stroke (44.70 ± 41.124) and STRd after cortico-striatal stroke (99.806 ± 0.434).

### Comparison between cortical and cortico-striatal stroke

Next, we directly compared the changes in SC-FC coupling between both stroke models in the sensorimotor regions. For descriptive decomposition of the model-specific pattern, regions were stratified according to the direction of the PT–MCAO ΔSC-FC coupling difference into PT-higher and MCAO-higher groups (***SI Appendix,* Table S2**). Mouse-level ΔSC-FC coupling was then averaged across regions within each directional group, and regional comparisons were subsequently used to identify the regions contributing most strongly to each pattern. After sorting, only contralateral to ischemic SC-FC coupling changes were significant after correction for multiple comparisons (**Fig. 3**).

**Figure 3:**
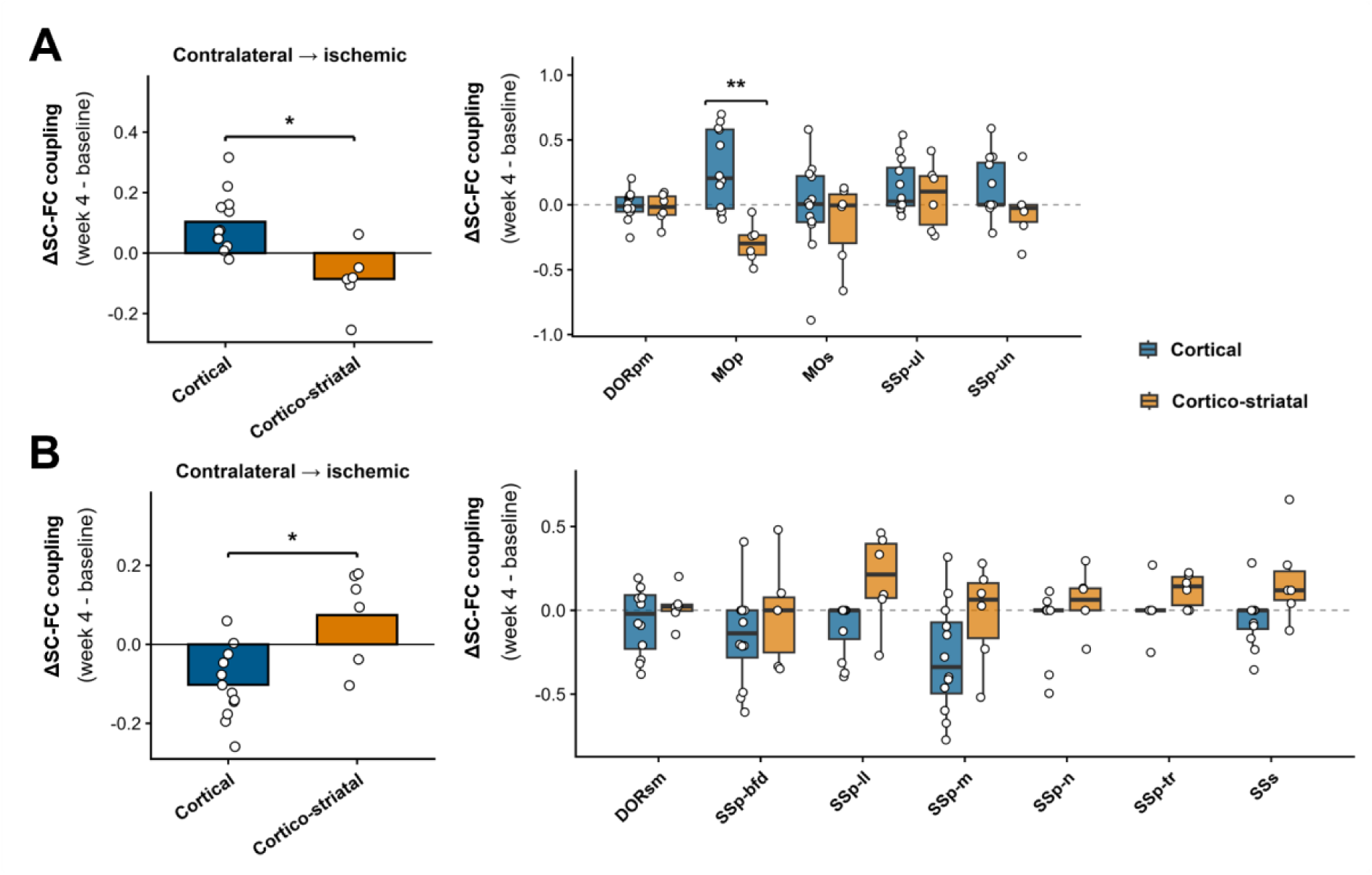
Contralateral-to-ischemic SC-FC coupling changes distinguish cortical from cortico-striatal stroke. Changes in coupling (ΔSC-FC; week 4 minus baseline) were compared between cortical and cortico-striatal stroke for contralateral-to-ischemic connections. (A) Regions with greater SC-FC coupling changes after cortical stroke showed a significant group-level difference between stroke models, driven predominantly by primary motor cortex (MOp). Regional ΔSC-FC coupling values are shown for DORpm, MOp, MOs, SSp-ul, and SSp-un. (B) Regions showing lower ΔSC-FC coupling changes in cortical compared with cortico-striatal stroke showed a significant group-level difference, although no individual regional comparison remained significant after FDR correction. Regional ΔSC-FC coupling values are shown for DORsm, SSp-bfd, SSp-ll, SSp-m, SSp-n, SSp-tr, and SSs. Bars in the left panels show mean group-level ΔSC-FC coupling, with individual animals overlaid. Box plots in the right panels show regional distributions. Bars show mean mouse-level ΔSC-FC coupling averaged across regions within each directional group; box plots show regional distributions (median and interquartile range), with circles indicating individual mice. Positive and negative values indicate increased and decreased coupling relative to baseline, respectively. Group-level and regional comparisons used independent-samples Student’s *t*-tests with Benjamini–Hochberg FDR correction across grouped comparisons and separately within each directional group, respectively. *FDR-adjusted P < 0.05; **P < 0.01.

In the group with higher SC-FC coupling changes in cortical compared with cortico-striatal stroke, the group-level difference was significant after FDR correction (adjusted P = 0.0110), with MOp showing the strongest regional difference and remaining significant after regional FDR correction (adjusted P = 0.0031; **Fig. 3A**). For contralateral-to-ischemic SC-FC coupling changes with lower values in cortical compared with cortico-striatal stroke, the group-level difference was significant after FDR correction (adjusted p = 0.011). The largest regional differences were observed in SSp-ll, SSp-tr, and SSs, although none remained significant after regional FDR correction (adjusted p = 0.093, 0.100, and 0.100, respectively; **Fig. 3B)**. The regional SC-FC coupling differences between stroke models could not simply be related to the difference in lesion size and only partly related to their distinct lesion distributions (***SI Appendix*, Fig. S6**). For contralateral-to-ischemic connections, MOp was the major contributor to higher coupling changes after cortical compared with cortico-striatal stroke (**Fig. 3A**). MOp was also significantly more affected by the cortical lesion than by MCAO (p < 0.0001), indicating that the distinct remodeling of contralateral MOp connectivity occurred in the context of different damage to its ipsilesional target network. However, this relationship was not consistent across sensorimotor regions. The regions contributing strongest to the opposite SC-FC coupling pattern in **Fig. 3B**, particularly SSp-ll, SSp-tr, and SSs, showed different degrees of lesion involvement and did not exhibit a uniform relationship between regional tissue damage and the direction or magnitude of SC-FC coupling change (***SI Appendix*, Fig. S6**).

Collectively, these findings indicate contralateral-to-ischemic SC-FC coupling as the primary circuit separating the two stroke models, with opposing motor- and sensorimotor-dominated patterns that may partly reflect differences in regional lesion distribution but did not show a simple correspondence with local tissue involvement (**Fig. 4A**).

**Figure 4:**
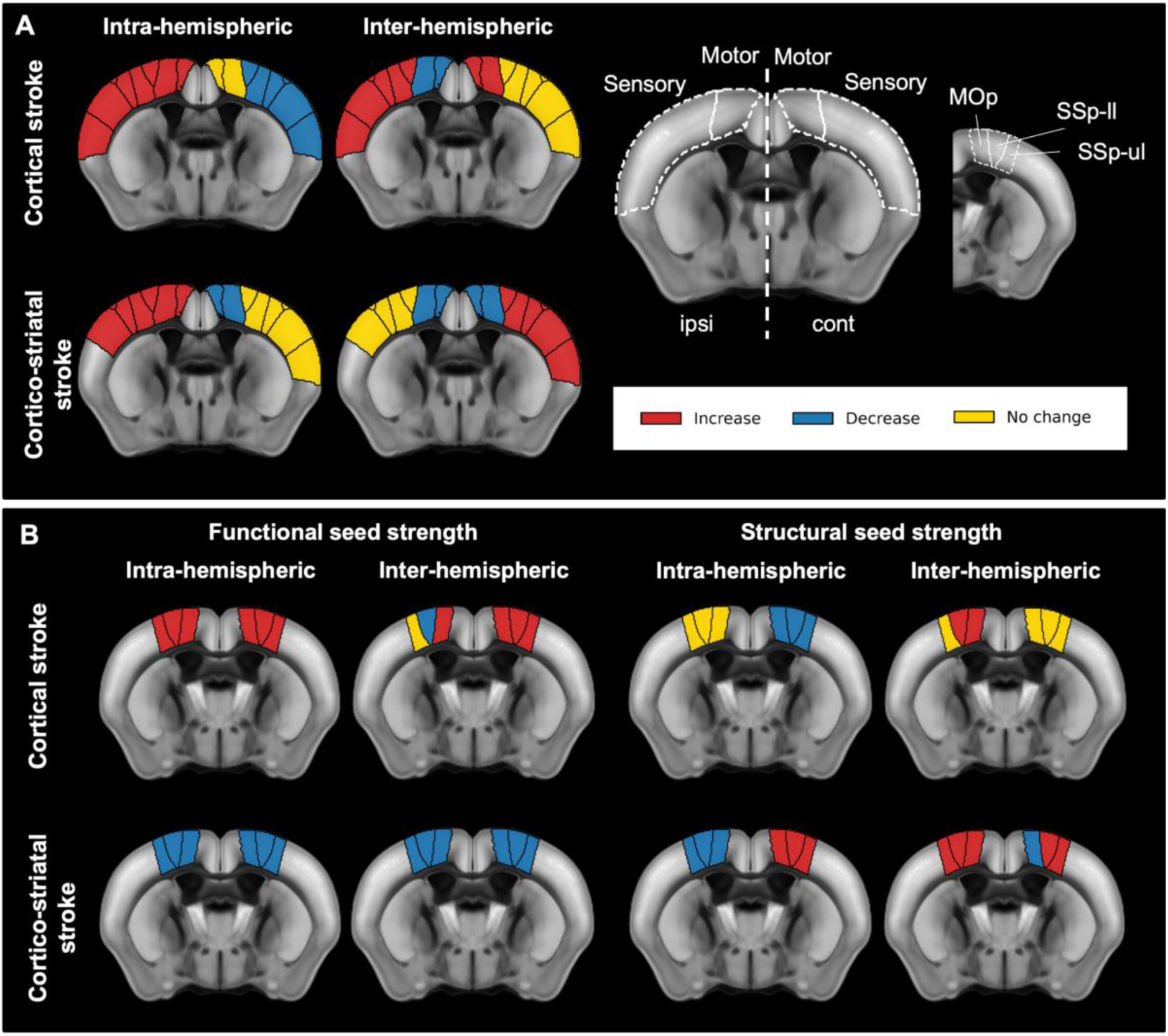
Distinct structure–function coupling patterns after cortical and cortico-striatal stroke. **(A)** Schematic summary of SC-FC coupling changes in representative motor and sensory regions at week 4. Intra-hemispheric SC-FC coupling changes within the ischemic hemisphere showed the same overall direction in both stroke models. Inter-hemispheric motor-region SC-FC coupling from the ischemic to the contralateral hemisphere likewise changed in the same direction. In contrast, SC-FC coupling changes within the contralateral hemisphere differed consistently between the two stroke models, and sensory-region SC-FC coupling from the ischemic to the contralateral hemisphere also showed opposing patterns. Labels: increased (red), decreased (blue), and no consistent change (yellow) compared to baseline. MOp, SSp-ll, and SSp-ul are shown on the Allen Mouse Brain Atlas template. **(B)** Corresponding functional (FSS) and structural seed-strength (SSS) changes. Within the ischemic hemisphere, cortical and cortico-striatal stroke showed opposing combinations of FSS and SSS changes despite yielding a similar net direction of SC-FC coupling change. In contrast, in the contralateral hemisphere, the two stroke models showed opposing FSS and SSS responses accompanied by different directions of SC-FC coupling remodeling. Interhemispheric FSS and SSS changes further illustrate that similar or divergent SC-FC coupling responses can arise from different combinations of structural and functional network alterations.

### Relationship between structural and functional network changes: consequences for structure-function coupling patterns

Next, we analyzed the underlying drivers of these complex coupling changes by comparing the separate roles of the structural and functional networks in the detected SC-FC coupling changes after stroke (**Fig. 5**). The analysis focused on MOp and SSp-ul/ll, as representative markers of sensorimotor deficits in both stroke models (18, 19). When comparing SC-FC coupling with structural and functional seed strength changes over time, we detected substantial differences between the stroke models and the four hemispheric subnetworks (**Fig. 5**). In the cortical stroke model, all three representative regions showed an increased intra-hemispheric functional seed strength (FSS) during weeks 1–2, whereas structural seed strength (SSS) remained largely unchanged within the ischemic hemisphere but decreased in the contralateral sensorimotor cortex. Inter-hemispheric connections originating from the contralateral hemisphere largely mirrored the contralateral intra-hemispheric pattern. In contrast, connections originating from the ischemic hemisphere displayed region-specific remodeling: MOp showed persistent increases in both FSS and SSS, SSp-ll showed increased SSS accompanied by reduced FSS, whereas SSp-ul remained structurally stable with reduced functional connectivity only at week 4.

**Figure 5:**
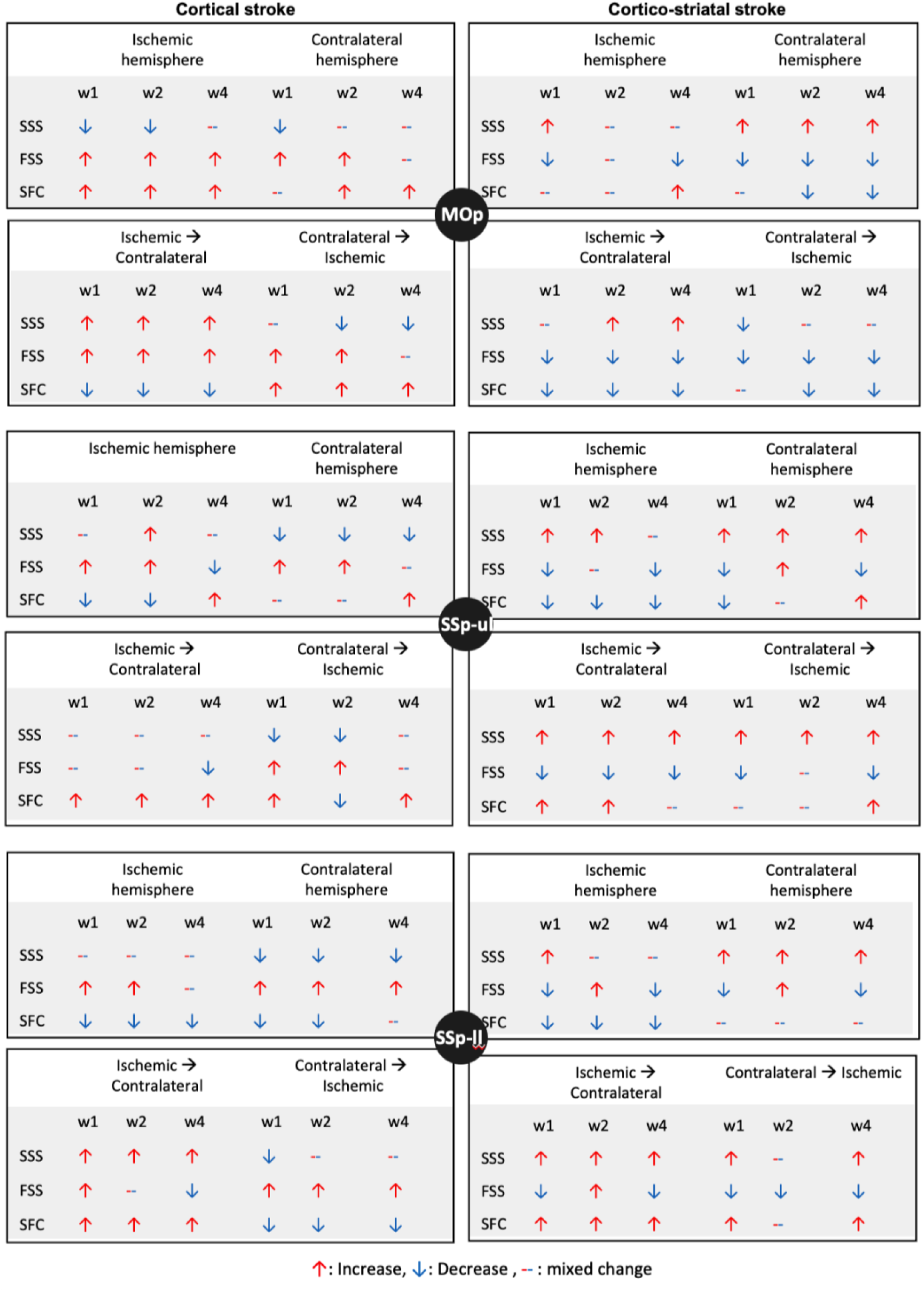
Summary of structural, functional and structure-function coupling changes after cortical and cortico-striatal stroke. Relative changes in structural seed strength (SSS), functional seed strength (FSS), and structure–function coupling (SFC) are shown for the primary motor cortex (MOp) and primary sensorimotor regions (SSp-ul and SSp-ll). Data are presented for the cortical stroke (left) and cortico-striatal stroke model (right), separately for intra-hemispheric connections within the ischemic and contralateral hemispheres and inter-hemispheric connections in both directions. Changes are shown relative to baseline at 1, 2, and 4 weeks after stroke. Upward red arrows indicate an overall increase, downward blue arrows an overall decrease, and horizontal red–blue bars mixed changes in seed strength relative to baseline. Arrows summarize the direction of observed changes and do not indicate statistical significance; statistical support for corresponding SC-FC coupling changes is provided in Figs. 2 and 3 and SI Appendix, Figs. S4 and S5.

In contrast, cortico-striatal stroke was characterized by reduced intra-hemispheric FSS in the ischemic hemisphere, while SSS shifted from early increases toward later decreases. In the contralateral hemisphere, SSS remained consistently elevated, whereas FSS was reduced or changed only modestly. Inter-hemispheric sensorimotor connections showed persistent structural increases, with predominantly reduced FSS in both directions. MOp showed sustained FSS reductions, along with increased SSS within the ischemic hemisphere and an early contralateral structural decrease.

Summarizing the trend of the seed strength changes across the four subnetworks over time, cortical and cortico-striatal stroke showed a widely opposing pattern of structural and functional remodeling (**Fig. 4B**). Despite these differences, distinct combinations of SSS and FSS changes after stroke converged on similar SC-FC coupling values within the ischemic hemisphere, particularly for motor cortex and intra-hemispherically for sensorimotor regions. In contrast, the more divergent contralateral seed-strength patterns paralleled the stroke model-specific SC-FC coupling differences. Thus, structural and functional seed-strength changes alone did not consistently distinguish the two stroke models, whereas their integration at the level of SC-FC coupling revealed the clearest model-specific network signatures.

## DISCUSSION

We provide a longitudinal characterization of SC-FC coupling up to 4 weeks post-stroke and compare two stroke models with largely different lesion extent and topology. In the healthy mouse brain, coupling was laterized in the SMN and showed opposite intra- vs. inter-hemispheric organization in SMN and DMN. Following stroke, coupling remodeling was dynamic, region- and direction-dependent, and extended beyond the ischemic territory as defined in the subacute phase. Although structural and functional seed-strength changes differed substantially between the two stroke models, SC-FC coupling changes within the ischemic hemisphere were comparatively similar, whereas the strongest stroke model-specific difference was detected from contralateral and contralateral-to-ischemic connections. The absence of a simple relationship between SC-FC coupling changes and local lesion burden further indicates that coupling captures distributed network reorganization, especially different roles of the intra- and inter-hemispheric network connections, rather than lesion effects alone.

### SC-FC coupling in the healthy brain

Higher SC-FC coupling in the SMN compared to the DMN is in full accord with the established sensorimotor-to-association gradient of SC-FC coupling in the human brain, together with evidence of decreasing coupling from unimodal cortex preferring local communication toward transmodal cortex preferring global communication. This gradient has been linked to the comparatively tight anatomical constraints on functional communication in sensory and motor networks, whereas functional interactions in transmodal systems, here the DMN, depend more strongly on distributed and polysynaptic network communication (4, 6, 20, 21). We further discovered a novel lateralization of SC-FC coupling in the mouse brain SMN, with stronger intra-hemispheric coupling in the right than in the left hemisphere. Importantly, this asymmetry was not evident in directional inter-hemispheric coupling, suggesting it primarily reflects differences in the internal organization of the left vs. right SMN rather than a general directional bias in inter-hemispheric communication. This result adds a novel component to the functional and structural lateralization of the human brain, including SMN (22, 23), which was only recently also described in rodents (24). In contrast, the DMN showed no significant left–right difference in intra-hemispheric SC-FC coupling, although equivalence between hemispheres was not formally established. This comparatively symmetric organization is consistent with the bilateral organization of resting-state networks and distributed connectivity of DMN regions reported in rodents (24, 25).

A second organizational principle emerged in the separate SC-FC coupling analysis of intra- and inter-hemispheric components. Previous studies have examined global, network-level, and intra-versus interhemispheric structure-function relationships, but region-wise coupling within specific functional networks has received considerably less attention. Global values ranged between 0.18 and 0.27 (7, 9, 26–28); 0.24 for visual and subcortical networks, and 0.14 for the DMN (5), as well as 0.3 for sensorimotor regions (29). Straathof et al. reported a substantially higher whole-cortex correlation in healthy rats (ρ = 0.41), which increased to ρ = 0.51 when restricted to intra-hemispheric connections, compared with ρ = 0.41 for inter-hemispheric connections (30). Although these values cannot be directly compared with our region-wise SC-FC coupling coefficients because of differences in analytical scale and correlation approach, their magnitude is remarkably similar to the coupling values observed in the healthy mouse brain here. Extending these observations, our region- and network-resolved analysis revealed a different situation of intra-hemispheric and inter-hemispheric SC-FC coupling values for the SMN and DMN. The inter-hemispheric SC-FC coupling of the SMN was substantially lower than intra-hemispherically (0.33 vs. 0.52). In contrast, for the DMN, the difference was reversed, i.e., the inter-hemispheric SC-FC coupling was higher than the intra-hemispheric coupling (0.48 vs. 0.31). This difference was also evident at the regional level. Most primary sensorimotor regions showed higher intra- than inter-hemispheric coupling. This organization is consistent with the strong topographic and reciprocal connectivity of primary sensorimotor and motor areas, in which function is closely linked to anatomically defined sensorimotor circuits (31, 32). This predominantly intrahemispheric organization is supported by Straathof et al., who found that 88% of cortical connections characterized by both strong diffusion-based SC and strong FC were intrahemispheric, with 62% of these connections located within the sensorimotor network (30). Interestingly, this intra-hemispheric dominance was absent in the two higher-order sensorimotor regions, SSs and MOs. Both regions are less restricted to primary sensorimotor processing and participate in more distributed cortical networks. MOs has extensive long-range cortical and subcortical projections (33), whereas SSs participates in extensive cross-regional and callosal sensorimotor connectivity (32). Thus, the similar intra-and inter-hemispheric coupling of SSs and MOs may reflect more balanced integration of local sensorimotor and distributed bilateral connectivity than in the more topographically organized primary sensorimotor regions.

The opposite pattern in the DMN is consistent with its distributed bilateral organization and long-range connectivity (24, 25, 34). In particular, ACA has direct callosal connections with its contralateral homolog, while RSP is a highly interconnected core region of the mouse DMN (34, 35). This distributed bilateral organization may explain why regions such as ACA, RSP and STRd showed higher inter- than intra-hemispheric SC-FC coupling. Importantly, higher inter-hemispheric coupling does not imply stronger inter-hemispheric connectivity itself, but rather indicates closer correspondence between structural and functional connectivity profiles across hemispheres. Thus, the opposite intra-versus inter-hemispheric coupling patterns of SMN and DMN may reflect different organizational principles: stronger local and topographic structural constraints in primary sensorimotor networks versus stronger bilateral and distributed integration in the DMN.

### SC-FC coupling changes after stroke

After stroke, SC–FC coupling remodeling was more pronounced in the SMN than the DMN and varied by region, hemisphere, direction, and time. Importantly, these changes showed no simple correspondence with regional lesion involvement: coupling remained stable in some substantially affected regions but changed in anatomically intact or minimally affected regions. This pattern is consistent with the concept of stroke as a network disorder, in which focal injury causes structural and functional disturbances extending into anatomically intact and contralateral regions (36–38). The dissociation between local tissue damage and SC-FC coupling changes was particularly evident in the DMN, where coupling remained comparatively stable despite substantial lesion involvement of individual regions (e.g., ACA, STRd, and TEa). This network-specific response may reflect the different structural-functional architectures of the SMN and DMN (4, 6, 21), with distributed organization of transmodal networks being less dependent on individual structural pathways and potentially providing greater resilience of SC-FC coupling. Conversely, pronounced but bidirectional SC-FC changes in the SMN may be particularly relevant to motor outcomes. Reduced dynamic SC-FC coupling has been associated with greater motor impairment after subcortical stroke (8), whereas stronger pathological motor-network coupling has also been associated with poorer motor function in acute stroke (39). Thus, an increase or decrease in SC-FC coupling cannot itself be interpreted as adaptive or maladaptive, but likely depends on the affected network, connection direction, and stage after stroke.

### SC-FC coupling changes depending on stroke model

The two stroke models differed most clearly in contralateral-to-ischemic coupling, with MOp contributing most strongly to the PT-higher pattern and SSp-ll, SSp-tr, and SSs to the opposite pattern. The prominent contribution of MOp is consistent with recruitment of the intact contralesional motor cortex after unilateral stroke, particularly when ipsilesional motor networks are compromised (36, 40). More recent work further indicates that the contribution of contralesional MOp depends on the structural reserve of the affected hemisphere (41). In experimental stroke, structural remodeling is not restricted to the lesioned cortex or to transcallosal pathways. The spared motor hemisphere undergoes dendritic and synaptic remodeling, while neurons in contralesional MOp can additionally sprout corticospinal projections toward denervated spinal targets (42). Such remodeling provides a plausible network context for the strong contralateral MOp contribution observed after cortical stroke. In the larger cortico-striatal lesion, however, disruption extends beyond cortex into subcortical motor circuits, potentially constraining or redirecting this inter-hemispheric cortical reorganization.

In contrast, the cortico-striatal stroke model involved the striatum and other subcortical tissue, disrupting the cortico-striato-thalamo-cortical circuit that contributes to motor-network reorganization (43). Regional lesion distribution may therefore contribute to the model-specific SC-FC coupling pattern, but the inconsistent relationship between lesion involvement and coupling across sensory regions indicates that lesion topology alone cannot account for it and again points to a stroke-induced network effect (36). This regional and hemispheric specificity extends early clinical observations of predominantly global poststroke decoupling (9). More recent clinical studies similarly indicate that SC-FC coupling abnormalities depend on lesion topology (7), differ between regions and hemispheres (10), and involve inter-hemispheric motor-network connections (10, 39). Thus, our finding that contralateral-to-ischemic rather than ipsilesional coupling most clearly separates the two stroke models supports an emerging view that poststroke SC-FC coupling is determined by the spatial organization of network reorganization rather than by a uniform loss of structural-functional correspondence. The functional relevance of these coupling patterns cannot be determined from the present analysis because SC-FC coupling was not directly related to behavioral outcome. Nevertheless, clinical studies showing associations between SC-FC coupling and motor impairment or recovery (9, 44) suggest that the region- and direction-specific patterns identified here warrant direct testing against longitudinal motor recovery.

### Decomposition of structural and functional components of SC-FC coupling

Decomposition into structural and functional seed strengths revealed that the direction of SC-FC coupling changes could not be inferred systematically from either measure. Depending on region, hemisphere, and connection direction, SC-FC coupling changes paralleled changes in SSS or FSS, but could also occur in the opposite direction despite concordant changes in both measures. Such dissociation between structural and functional connection strength is already evident in the healthy rodent and human brain (30, 45). This apparent dissociation reflects the different information captured by these metrics: SSS and FSS quantify mean structural and functional connection strength, whereas SC-FC coupling quantifies the correspondence between their connectivity profiles across target regions. Thus, simultaneous increases in SSS and FSS can be accompanied by reduced SC-FC coupling when structural and functional remodeling affects different connections within these profiles. This distinction is particularly relevant for comparing the two stroke models and may explain why SC or FC, considered separately, showed less distinct separation between cortical and cortico-striatal stroke in our previous analyses (12, 13). The decomposition therefore shows that SC-FC coupling is neither a surrogate for SC nor FC, but provides complementary information about how functional network organization relates to the underlying structural network. Such divergence between structural and functional remodeling is consistent with longitudinal stroke studies showing that structural and functional connectivity can evolve along different trajectories and differ in their relationship to motor recovery (46). Structural disruption can constrain functional connectivity, particularly across inter-hemispheric pathways, but this relationship is not one-to-one and varies across networks and pathways. Our results extend this principle by showing that divergence between mean structural and functional strength does not itself predict the direction of SC-FC coupling change.

Similar coupling values can therefore arise from substantially different structural and functional network states and should not be interpreted as evidence of network preservation. Rather, SC–FC coupling may capture the evolving correspondence between the structural network remaining after injury and the functional network reorganizing on this substrate. Poststroke processes including axonal and synaptic remodeling and vascular plasticity could contribute to this evolving network organization during the weeks after injury (17, 18, 47), although these mechanisms were not directly assessed here. Consistent with this interpretation, clinical studies indicate that poststroke SC-FC coupling and its relationship with motor function depend on network, pathway, time after stroke, impairment severity, and recovery trajectory (8, 44, 48).

### Methodological limitations

A methodological limitation of the present study is that the cortical and cortico-striatal stroke datasets originated from previously established longitudinal MRI studies with cohorts of unequal sample size, mouse strain, lesion characteristics, and acquisition protocols. Cohort- or acquisition-specific factors may have contributed to the between-model differences observed after stroke. In particular, the smaller cortico-striatal cohort may have reduced statistical power to detect subtle regional effects. The two stroke models differed not only in lesion extent and topology but also in their underlying ischemic pathophysiology. Photothrombosis produces a well-defined cortical lesion through local small-vessel thrombosis and is characterized by a comparatively limited ischemic border zone, whereas MCAO causes territorial ischemia that commonly extends into subcortical structures and, after filament withdrawal, includes a defined reperfusion phase (47, 49). These differences in vascular occlusion, peri-infarct tissue characteristics, and reperfusion can influence the subsequent evolution of tissue injury and recovery. Moreover, SC-FC coupling represents an indirect relationship between structural connectivity estimated from diffusion MRI tractography and functional connectivity derived from resting-state BOLD correlations; therefore, changes in coupling should not be interpreted as direct evidence of underlying axonal or synaptic mechanisms. Finally, all experiments were performed in male mice and behavioral assessments were not sufficiently harmonized across cohorts to directly relate SC-FC coupling changes to behavioral recovery.

## DATA, MATERIALS, AND SOFTWARE AVAILABILITY

MRI data are available from GIN (DTI: https://doi.org/10.12751/g-node.ln5k5d and rs-fMRI: https://doi.org/10.12751/g-node.pmvtz1). The AIDAmri software is available open-access on GitHub (https://github.com/Aswendt-Lab/AIDAmri), and the custom R and Python codes used for the analyses are publicly accessible (https://doi.org/10.12751/g-node.rn9kwg).

## ACKNOWLEDGEMENTS

We are grateful for the financial support by grants from the Deutsche Forschungsgemeinschaft (DFG, German Research Foundation): project ID 431549029–SFB 1451 and the Henry-Oswalt Foundation.

