## Supplementary material for "Structure-function coupling reveals hemisphere-specific network reorganization after stroke": SI Appendix

### Supporting Information

#### Stroke models

Permanent cortical stroke was induced by photothrombosis as described previously (1). Briefly, we administered Rose Bengal as a photosensitive dye, and targeted cortical illumination caused localized endothelial damage and thrombus formation, resulting in a small cortical infarct. Transient cortico-striatal stroke was induced by MCAO as described previously (2). Briefly, a coated intraluminal filament was inserted into the internal carotid artery for 30 min and then withdrawn to allow reperfusion, producing a larger cortico-striatal lesion.

#### MRI acquisition

MRI was performed on a 9.4 T Bruker BioSpec 94/20USR system (Bruker BioSpin, Ettlingen, Germany) equipped with a cryogenic coil and operated with ParaVision v6.0.1. We anesthetized mice with isoflurane (2-3% in 70/30 N2/O2), positioned them in an animal carrier/cradle, and monitored respiration and body temperature throughout scanning to maintain physiological stability. High-resolution T2-weighted images were acquired in all animals before diffusion MRI or resting-state fMRI (***SI Appendix*, Table S1**).

#### Image preprocessing and atlas registration

All MRI data were processed with AIDAmri version 1.3 (3) and registered to the Allen Mouse Brain Atlas (AMBA; Common Coordinate Framework version 3.0) (4). Shared preprocessing steps included conversion from Bruker raw data to NIfTI format, brain extraction with FSL BET, and correction of bias field inhomogeneities using the Multiplicative Intrinsic Component Optimization (MICO) algorithm (4). Anatomical T2-weighted images were aligned to atlas space using affine and non-linear registration. Modality-specific data were then coregistered to the corresponding anatomical scans for region-based analysis.

For structural connectivity, the diffusion data were registered to the T2-weighted anatomical scans before tractography and matrix generation. For functional connectivity, additional preprocessing steps included slice-timing correction, motion correction, regression of respiratory-related fluctuations, temporal band-pass filtering between 0.01 and 0.1 Hz, and spatial smoothing with a Gaussian kernel of sigma 0.1 mm. After co-registration to the anatomical images and transformation to atlas space, regional time series were extracted from 98 atlas-defined regions.

#### Lesion segmentation

Ischemic lesions were identified as hyperintense regions on T2-weighted MRI and segmented semi-automatically using ITK-SNAP with the 3D snake / active contour tool (5). The resulting stroke masks were transformed to atlas space, enabling generation of lesion incidence maps and region-specific lesion quantification. This approach allowed both qualitative comparison of lesion topology and quantitative evaluation of lesion involvement across the two stroke models.

#### Lesion masking and exclusion strategy

Because post-stroke edema and infarct-related signal abnormalities can distort both diffusion MRI and resting-state fMRI, lesion-aware masking procedures were applied in both modalities. In both workflows, lesion masks generated at week 1 were registered to the corresponding scans at later time points.

For structural connectivity, the transformed lesion areas were set to zero in the fiber orientation distribution maps before tractography. Fiber-count matrices were subsequently recalculated, and the volumes of regions partially affected by lesion exclusion were updated so that corrected region sizes could be used for calculation of fiber-density matrices. This strategy was intended to reduce distortion of diffusion-based structural connectivity estimates caused by edema or infarct tissue.

For functional connectivity, we excluded lesion voxels from regional time-series extraction by setting transformed lesion areas to zero in the time-series maps while retaining their spatial positions as zero-valued placeholders in the connectivity matrices. Anatomical segmentations and rs-fMRI data were therefore modified such that only intact tissue contributed to regional signal averaging, while the spatial organization of the network matrices was preserved.

#### Structural connectivity analysis

We performed whole-brain tractography and structural connectivity analysis in DSI Studio (6), version 2023, as reported in the structural connectivity dataset. Deterministic fiber tracking was conducted across the whole brain using a modified AMBA template comprising 98 selected regions of interest. Tracking parameters included a fiber length threshold of >0.5 mm and <120 mm and a turning angle of <55°. For each animal and time point, the tractography output was converted into an adjacency matrix in which each entry represented the number of fibers connecting two brain regions. Connection weights were transformed into fiber density (FD) following Hagmann et al. (7), with modifications to account for region size and fiber length, reducing potential bias toward larger regions or longer tracts. Seed strength for SC was defined as the average fiber density of a given seed region relative to other regions within a specified matrix compartment. Using the fiber-density matrices, intra-hemispheric and inter-hemispheric connections were extracted separately and averaged for each region to quantify structural connectivity changes over time in the ischemic hemisphere, contralateral hemisphere, and between hemispheres.

#### Functional connectivity analysis

For each of the 98 atlas-defined brain regions, a representative resting-state fMRI time series was obtained by averaging the voxel-wise signal within that region after lesion exclusion. Pairwise Pearson correlation coefficients were computed between all regional time series to generate full FC matrices. Correlation coefficients were transformed to Fisher-z scores to enable averaging and statistical analysis.

Seed strength for FC was defined as the mean Fisher-z-transformed FC between a given seed region and all other regions within a specified matrix compartment. Intra-hemispheric and inter-hemispheric seed strengths were calculated separately, mirroring the hemispheric compartmentalization used in the structural connectivity analysis.

#### Table S1: MRI parameters

| Study | Sequence | TR (ms) | TE (ms) | Voxel (µm$^3$) | FOV (mm$^2$) | B-value (s/mm$^2$) | Gradient directions |
|---|---|---|---|---|---|---|---|
| PT | Turbo-RARE | 5500 | 32.5 | 68.4 × 68.4 × 300 | 17.5 × 17.5 | NA | NA |
| PT | DTI | 3000 | 17.5 | 141 × 141 × 400 | 18 × 18 | 677 | 30 |
| PT | rs-fMRI | 1420 | 18 | 182 × 182 × 500 | 17.5 × 17.5 | NA | NA |
| MCAO | Turbo-RARE | 5500 | 32.5 | 68.4 × 68.4 × 200 | 17.5 × 17.5 | NA | NA |
| MCAO | DSI (8× Qball) | 3500 | 20 | 139 × 139 × 500 | 17.8 × 17.8 | 2000 | 126[1] |
| MCAO | rs-fMRI | 2840 | 18 | 182 × 182 × 500 | 17.5 × 17.5 | NA | NA |

[1] Half sphere. Flip angle (90°) and refocusing flip angle (180°) for all sequences. DSI: diffusion spectrum imaging; DTI: diffusion tensor imaging; FOV: field of view; MCAO: middle cerebral artery occlusion; NA: not available; PT: photothrombosis; rs-fMRI: resting-state functional magnetic resonance imaging; TE: echo time; TR: repetition time.

#### Table S2: SC-FC coupling values contralateral to ischemic comparison

Cortical = PT; Cortico-striatal = MCAO; p = raw Student t-test p-value; q = FDR corrected p-value

| Analysis level | Directional group | Region / summary | Cortical mean ± SD | Cortico-striatal mean ± SD | p-value | q-value |
|---|---|---|---|---|---|---|
| Group-level | Group 1 (PT-higher regions) | Averaged directional regions | 0.103 ± 0.098 | -0.086 ± 0.102 | 0.0015 | 0.0110 |
| Group-level | Group 2 (MCAO-higher regions) | | -0.102 ± 0.090 | 0.074 ± 0.118 | 0.0028 | 0.0110 |
| Regional | Group 1 | DORpm | -0.006 ± 0.114 | -0.026 ± 0.116 | 0.7296 | 0.7296 |
| | | MOp | 0.273 ± 0.306 | -0.295 ± 0.153 | 0.0006 | 0.0031 |
| | | MOs | -0.016 ± 0.362 | -0.137 ± 0.318 | 0.4980 | 0.6912 |
| | | SSp-ul | 0.135 ± 0.207 | 0.067 ± 0.260 | 0.5530 | 0.6912 |
| | | SSp-un | 0.130 ± 0.231 | -0.037 ± 0.246 | 0.1778 | 0.4444 |
| | Group 2 | DORsm | -0.060 ± 0.199 | 0.021 ± 0.111 | 0.3691 | 0.3691 |
| | | SSp-bfd | -0.159 ± 0.283 | -0.017 ± 0.308 | 0.3420 | 0.3691 |
| | | SSp-ll | -0.101 ± 0.162 | 0.184 ± 0.276 | 0.0133 | 0.0931 |
| | | SSp-m | -0.285 ± 0.330 | -0.026 ± 0.297 | 0.1256 | 0.2197 |
| | | SSp-n | -0.059 ± 0.183 | 0.054 ± 0.177 | 0.2280 | 0.3192 |
| | | SSp-tr | 0.002 ± 0.111 | 0.120 ± 0.100 | 0.0429 | 0.1000 |
| | | SSs | -0.054 ± 0.156 | 0.182 ± 0.267 | 0.0299 | 0.1000 |

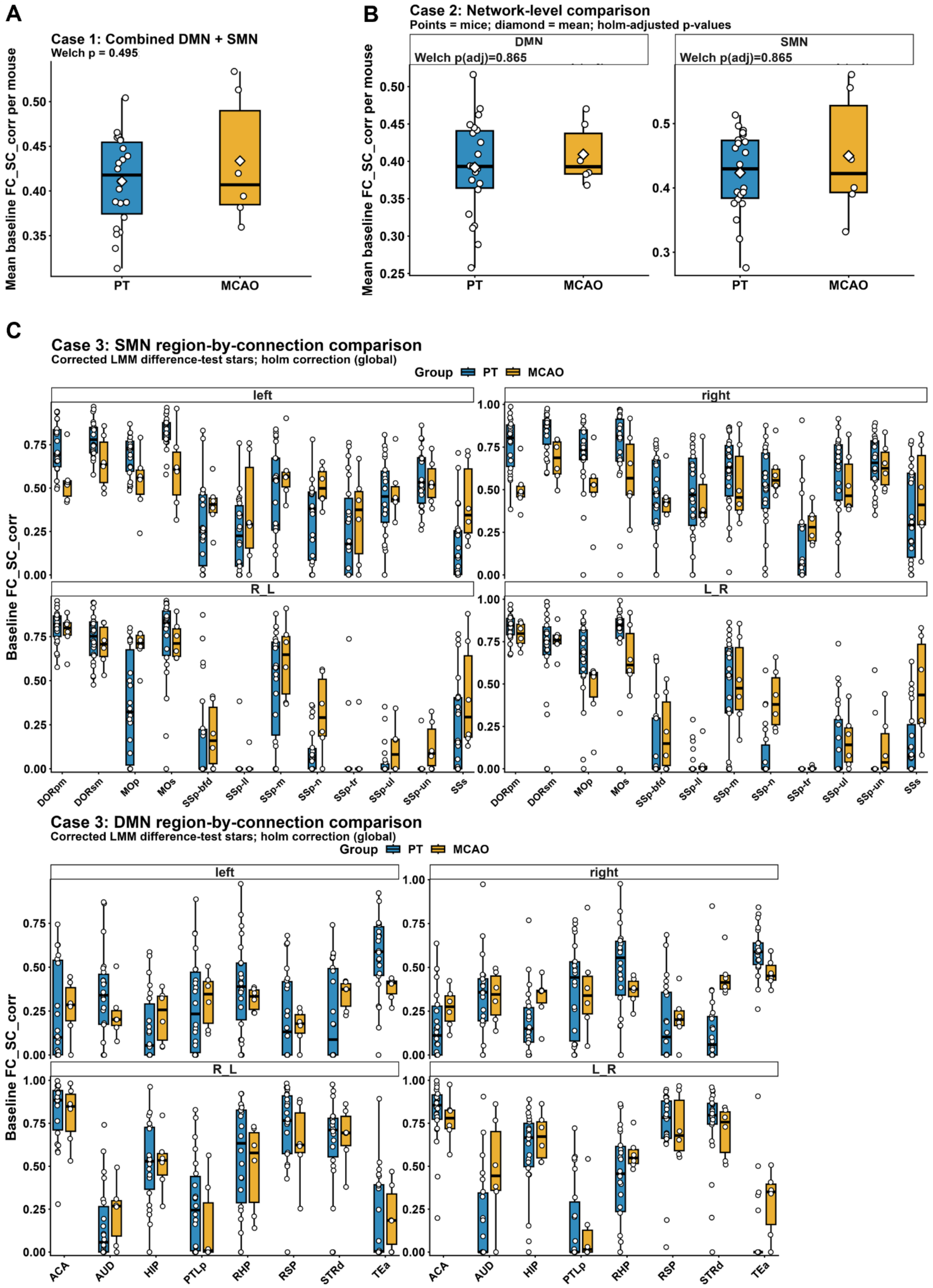


**Figure S1: Healthy brain baseline SC-FC coupling comparison between the PT and MCAO groups.** Baseline structure–function coupling (SC-FC coupling) was compared between mice assigned to the cortical stroke group (PT; n=22) and the cortico-striatal stroke group (MCAO; n=6) at three levels of analysis. For Cases 1 and 2, one mean SC-FC coupling value was calculated per mouse, and the independent groups were compared using two-sided Welch's t-tests. **A)** In Case 1, SC-FC coupling was averaged across the four connection categories and all 20 available DMN and

SMN regions for each mouse. Mean SC-FC coupling was 0.411±0.050 in the PT group and 0.434±0.073 in the MCAO group. The PT–MCAO difference was -0.0228 (95% CI [-0.0988,0.0532]), with no significant group difference according to Welch's (t)-test (P=0.495). **B)** In Case 2, PT and MCAO mice were compared separately within the DMN and SMN. The Welch t-test P-values were corrected across the two networks using the Holm method. In the DMN, mean SC-FC coupling was 0.392±0.064 for PT and 0.409±0.041 for MCAO, corresponding to a PT–MCAO difference of -0.0175 (95% CI [-0.0643,0.0293]). No significant group difference was detected (raw Welch P=0.432, Holm-adjusted P=0.865). In the SMN, mean SC-FC coupling was 0.424±0.063 for PT and 0.450±0.097 for MCAO, with a difference of -0.0263 (95% CI [-0.1279,0.0752]). The groups did not differ significantly (raw Welch P=0.551, Holm-adjusted P=0.865). Thus, the network-level results provided no evidence of a significant baseline group difference. **C)** In Case 3, regional and connection-specific SC-FC coupling values were analyzed using a linear mixed-effects model of the form SC-FC coupling~Group×Region×Connection+(1∣MouseID), with a random intercept for mouse and Satterthwaite approximation of the degrees of freedom. The model included 2,240 observations from 28 mice, 20 regions, and four connection categories: left and right intra-hemispheric coupling and the two directional inter-hemispheric connections (R_L) and (L_R). The omnibus analysis showed no overall main effect of group ($F_{1,26}$=0.809, P=0.377). Significant effects were found for region ($F_{19,2054}$=48.07, P=1.34×$10^{-148}$), connection category ($F_{3,2054}$=5.77, P=0.000628), and their interaction ($F_{57,2054}$=13.60, P=9.55×$10^{-105}$). The Group × Region interaction ($F_{19,2054}$=3.22, P=3.17×$10^{-6}$) and Group × Connection interaction ($F_{3,2054}$=2.66, P=0.0465) were significant, whereas the Group × Region × Connection interaction was not ($F_{57,2054}$=1.22, P=0.123). Model-adjusted PT–MCAO contrasts were subsequently calculated for each of the 80 region-by-connection combinations, with global Holm correction. No individual PT–MCAO contrast remained significant after correction; the smallest adjusted P-value was 0.0601. Consequently, the absence of significance stars in panel C indicates that no regional PT–MCAO difference survived multiplicity correction.
PT mice are shown in blue and MCAO mice in orange. Circles represent individual animals. Boxes show the median and interquartile range, with whiskers extending to 1.5 times the interquartile range. White diamonds in panels A and B indicate group means. Significance stars in panel C were based exclusively on globally Holm-adjusted linear mixed-model difference-test P-values. Collectively, these analyses detected no significant baseline difference between PT and MCAO mice at the combined or network level and no corrected region-by-connection differences.

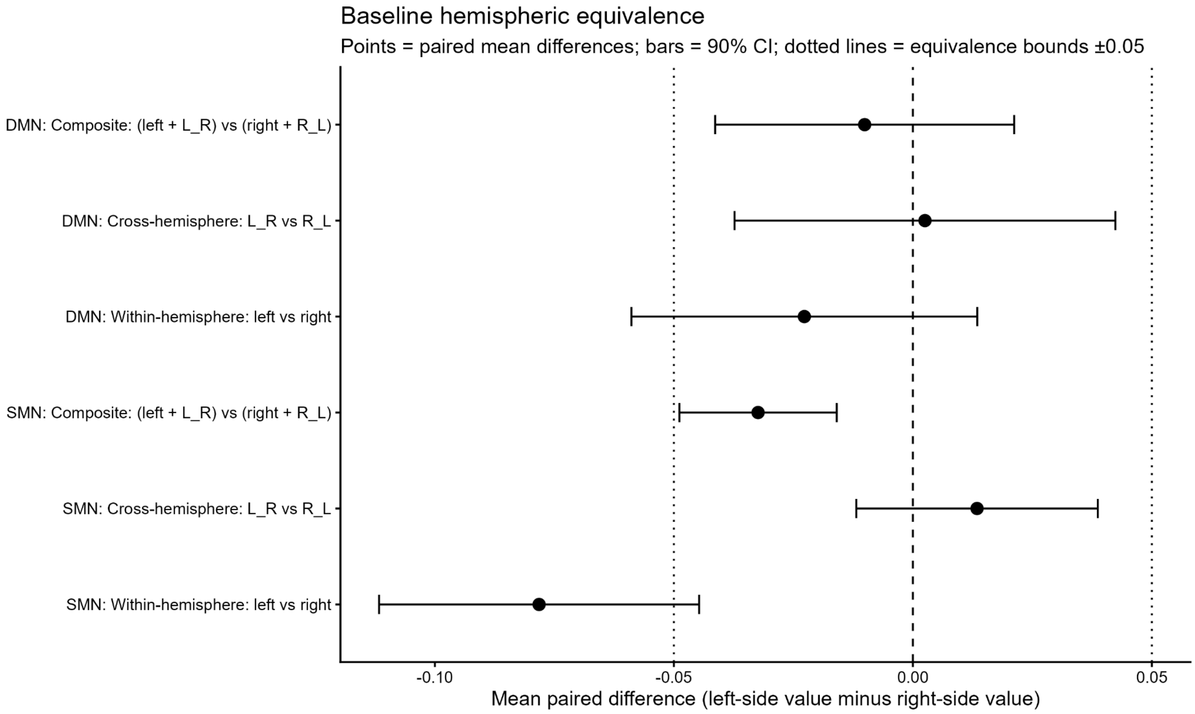


**Figure S2: Healthy baseline left-right hemisphere similarity of SC-FC coupling.** Baseline hemispheric similarity of SC-FC coupling was assessed in 28 mice pooled across the PT and MCAO groups using paired two one-sided tests (TOST) for equivalence. We detected no significant effect of the experimental group on paired hemispheric differences for any comparison (all $P \geq 0.095$). For each mouse, three paired comparisons were evaluated separately in the default mode network (DMN) and sensorimotor network (SMN): within-hemisphere coupling (left versus right), cross-hemisphere coupling (L_R versus R_L), and a composite hemispheric measure, calculated as the average of left and L_R values versus the average of right and R_L values. Points represent the mean paired difference, calculated as the left-side value minus the right-side value, and horizontal bars represent the corresponding 90% confidence intervals. The central dashed line indicates a difference of zero, and the dotted lines indicate the predefined equivalence bounds of ±0.05. Equivalence was concluded when the entire 90% confidence interval was contained within these bounds, and the TOST P-value was below 0.05. In the DMN, equivalence was established for the composite hemispheric measure (mean difference =−0.0101, 90% CI [−0.0414, 0.0212], TOST P=0.0193) and for cross-hemisphere coupling (mean difference =0.0025, 90% CI [−0.0373, 0.0424], TOST P=0.0262). Equivalence was not established for the direct left vs. right within-hemisphere comparison because its confidence interval extended beyond the lower equivalence bound (mean difference =−0.0227, 90% CI [−0.0589, 0.0135], TOST P=0.1048; however, an ordinary paired t-test also provided no evidence of a left–right difference (P=0.295). In the SMN, equivalence was established for the composite measure (mean difference =−0.0324, 90% CI [−0.0488, −0.0159], TOST P=0.0397) and for cross-hemisphere coupling (mean difference =0.0134, 90% CI [−0.0118, 0.0387], TOST P=0.0102). In contrast, the direct within-hemisphere comparison was not equivalent (mean difference =−0.0782, 90% CI [−0.1117, −0.0447], TOST P=0.9186) and showed significantly greater coupling in the right than in the left hemisphere in an ordinary paired t-test (P=0.00047). Thus, the composite and directional cross-hemisphere measures supported baseline hemispheric similarity in both networks within the predefined ±0.05 range, whereas direct within-hemisphere equivalence was not demonstrated.

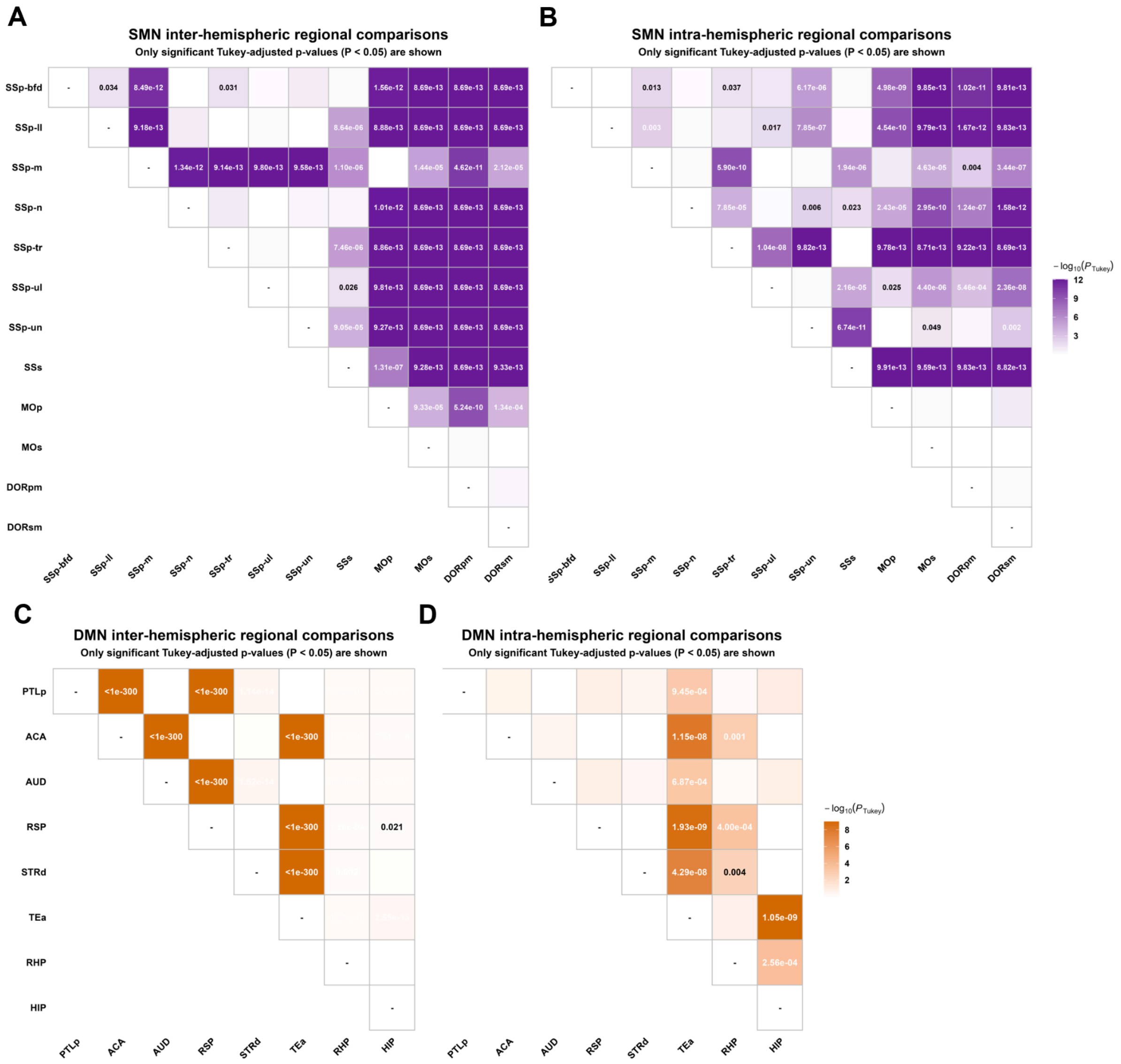


**Figure S3: Pooled regional pairwise comparisons of baseline structure–function coupling in the sensorimotor and default mode networks.** We analyzed baseline structure–function coupling (SC-FC coupling) jointly in 28 mice from the PT (n=22) and MCAO (n=6) groups, including the experimental group as a fixed effect. For each network and connection type, regional differences were assessed using a linear mixed-effects model of the form Coupling~Group+Region+(1∣SubjectID), adjusting the regional estimates for experimental group while accounting for repeated regional measurements within each mouse. Intra-hemispheric coupling was calculated for each mouse and region as the mean of the left- and right-hemisphere values, whereas inter-hemispheric coupling was calculated as the mean of the two directional cross-hemisphere values (L_R and R_L). Pairwise regional comparisons were performed on the estimated marginal means and corrected for multiple comparisons using the Tukey procedure. Heatmaps show pairwise comparisons for (A) SMN inter-hemispheric coupling, (B) SMN intra-hemispheric coupling, (C) DMN inter-hemispheric coupling, and (D) DMN intra-hemispheric coupling. Only significant Tukey-adjusted P-values (P<0.05) are displayed; blank off-diagonal cells indicate nonsignificant comparisons, and dashes identify comparisons of each region with itself. Cell shading represents −log10(PTukey), with darker colors indicating stronger statistical evidence. Extremely small adjusted P-values below the numerical display limit are reported as P<1×10−300. Overall, significant regional differences were more widespread for inter-hemispheric coupling, particularly in the DMN, whereas DMN intra-hemispheric differences were comparatively concentrated in comparisons involving TEa, RHP, and HIP.

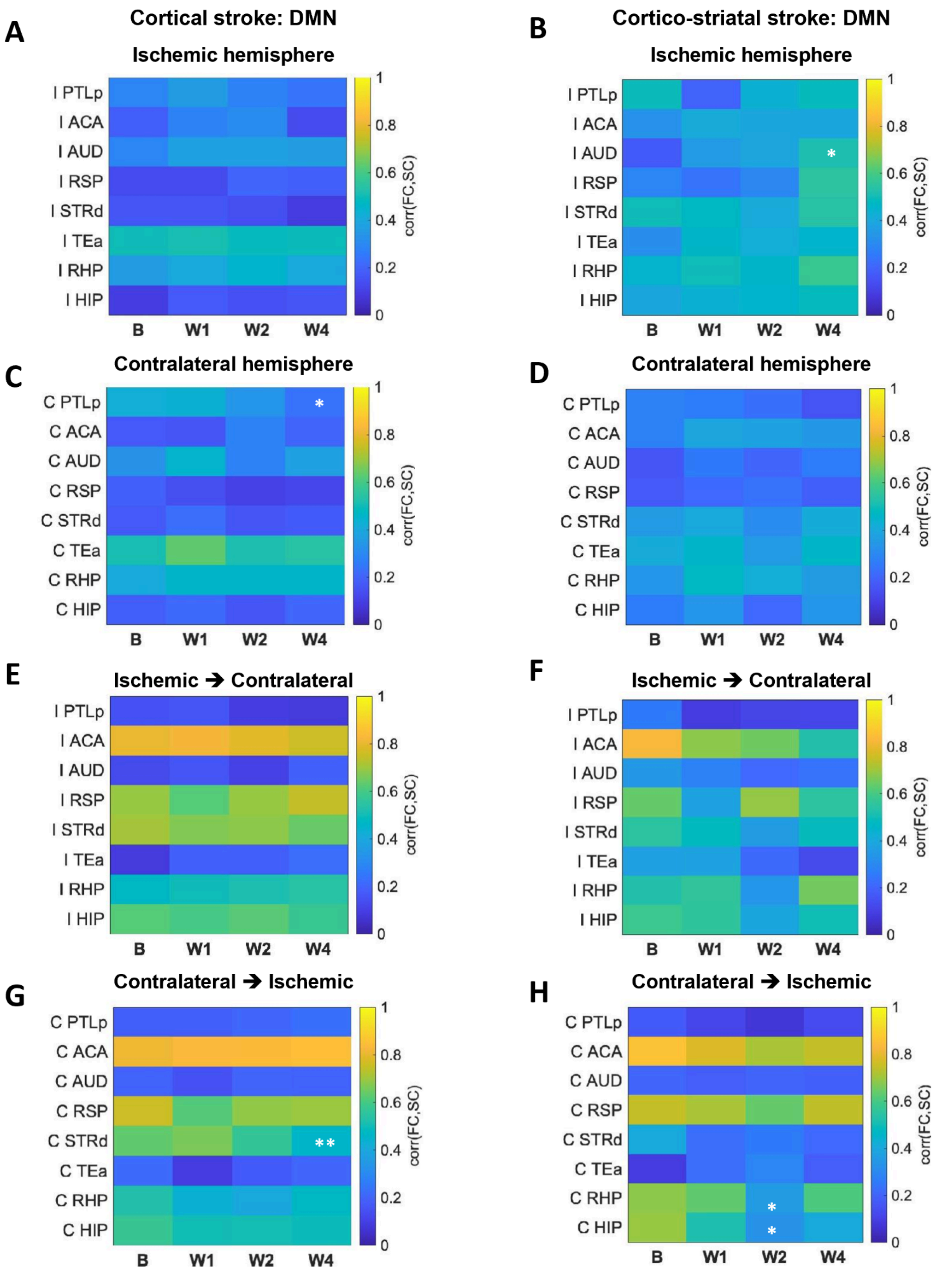


**Figure S4: Regional structure–function coupling (SC-FC coupling) changes in the default mode network after stroke.** A, C, E, G) SC-FC coupling values after cortical stroke, and B, D, F, H) SC-FC coupling values after cortico-striatal stroke. A, B) intra-hemispheric connections within the ischemic hemisphere; C, D) intra-hemispheric connections within the contralateral hemisphere; E, F) inter-hemispheric connections from the ischemic to the contralateral hemisphere; and G, H) inter-hemispheric connections from the contralateral to the ischemic hemisphere. SC-FC coupling is shown separately for each contributing seed region at baseline (B) and at 1, 2, and 4 weeks after stroke (W1, W2, and W4, respectively). Changes from baseline were assessed using linear mixed-effects models with Day, Region, and their interaction as fixed effects and MouseID as a random intercept (Coupling ~ Day × Region + (1 | MouseID)). Estimated marginal means were calculated for each day within each region, and W1, W2, and W4 were compared with baseline using treatment-versus-control contrasts. P values were corrected for multiple comparisons using the Benjamini–Hochberg false discovery rate (FDR) procedure. Significant changes from baseline were observed at W4 in the ischemic AUD for intra-hemispheric connections after cortico-striatal stroke (B), at W4 in the contralateral PTLp for intra-hemispheric connections after cortical stroke (C), at W4 in the contralateral STRd for inter-hemispheric connections from the contralateral to the ischemic hemisphere after cortical stroke (G), and at W2 in the contralateral RHP and HIP for inter-hemispheric connections from the contralateral to the ischemic hemisphere after cortico-striatal stroke (H). FDR-adjusted $p < 0.05$ (*) and $p < 0.01$ (**).

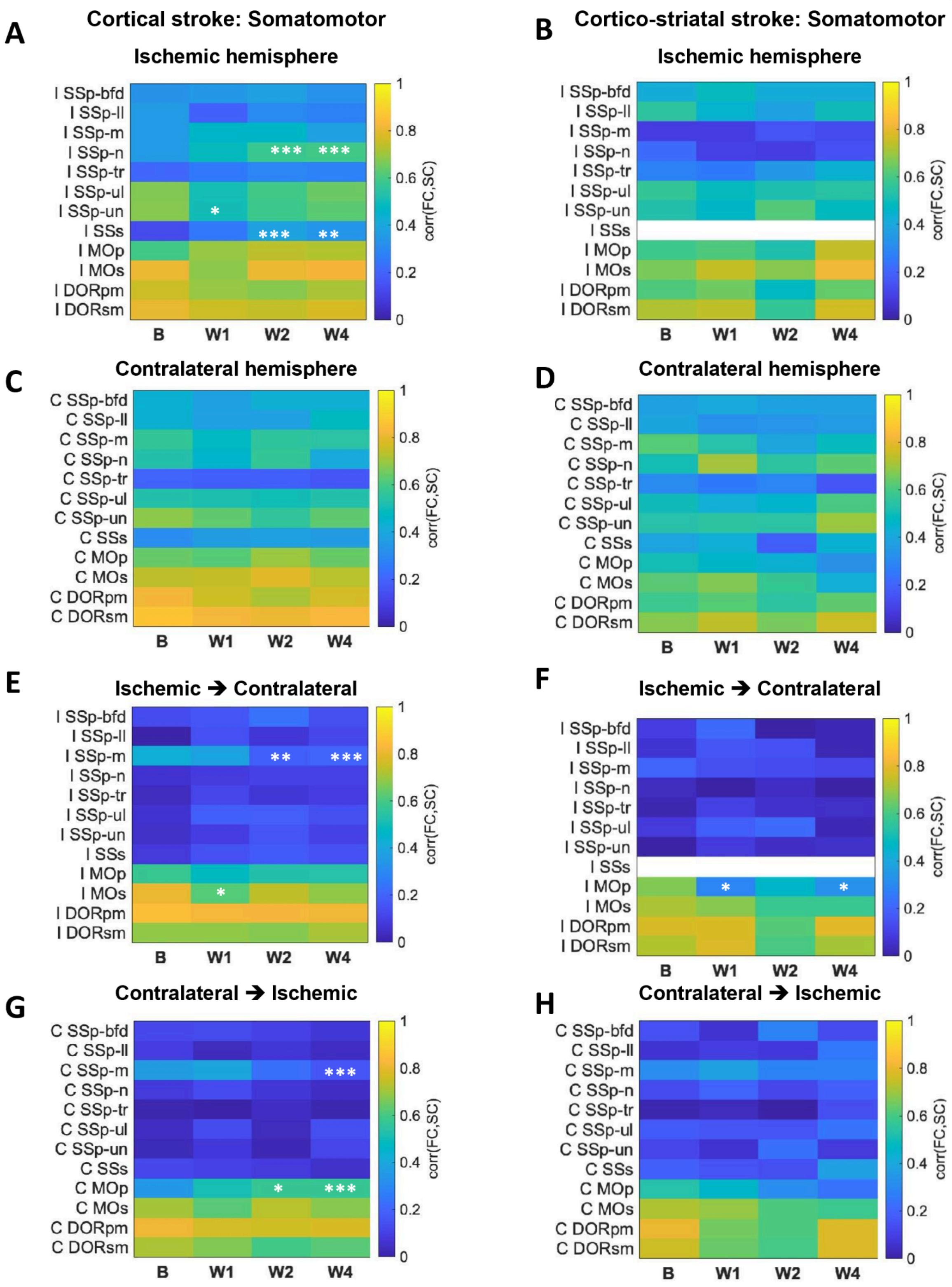


**Figure S5: Longitudinal SC-FC coupling changes in the sensorimotor network after stroke .** A, C, E, G) SC-FC coupling values after cortical stroke, and B, D, F, H) SC-FC coupling values after cortico-striatal stroke. A, B) intra-hemispheric connections within the ischemic hemisphere; C, D) intra-hemispheric connections within the contralateral hemisphere; E, F) inter-hemispheric connections from the ischemic to the contralateral hemisphere; and G, H) inter-hemispheric connections from the contralateral to the ischemic hemisphere. SC-FC coupling is shown separately for each contributing seed region at baseline (B) and at 1, 2, and 4 weeks after stroke (W1, W2, and W4, respectively). Changes from baseline were assessed using linear mixed-effects models with Day,

Region, and their interaction as fixed effects and MouseID as a random intercept (Coupling ~ Day × Region + (1 | MouseID)). Estimated marginal means were calculated for each day within each region, and W1, W2, and W4 were compared with baseline using treatment-versus-control contrasts. *P* values were corrected for multiple comparisons using the Benjamini–Hochberg false discovery rate (FDR) procedure. After cortical stroke, significant changes from baseline were observed for intra-hemispheric connections within the ischemic hemisphere (A) in SSp-n at W2 and W4, SSp-un at W1, and SSs at W2 and W4; for inter-hemispheric connections from the ischemic to the contralateral hemisphere (E) in SSp-m at W2 and W4 and MOs at W1; and for inter-hemispheric connections from the contralateral to the ischemic hemisphere (G) in SSp-m at W4 and MOp at W2 and W4. After cortico-striatal stroke, significant changes were observed for inter-hemispheric connections from the ischemic to the contralateral hemisphere (F) in MOp at W1 and W4. No significant changes from baseline were detected in panels C, B, D, or H. * indicates FDR-adjusted p < 0.05, ** indicates FDR-adjusted p < 0.01, and *** indicates FDR-adjusted p < 0.001.

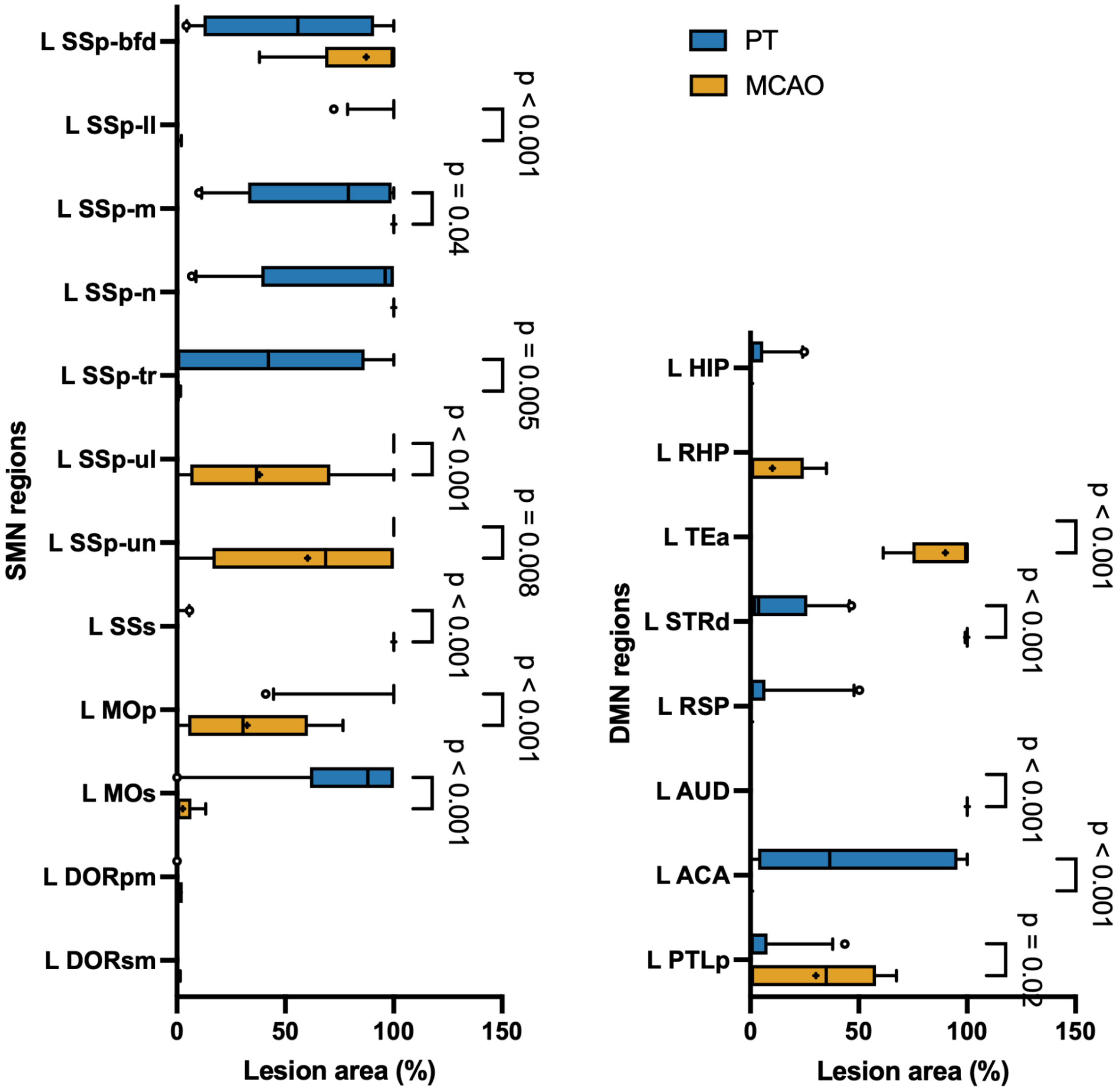


**Figure S6: Stroke lesion area per brain region in SMN and DMN at 1 wk after cortical vs. cortico-striatal stroke measured by T2-weighted MRI.** Lesion area was quantified at 1 wk after stroke using T2-weighted magnetic resonance imaging (MRI) and expressed as a percentage of the corresponding Allen Mouse Brain atlas brain region. Lesion area is shown for individual sensorimotor network (SMN; left) and default mode network (DMN; right) regions in the peri-lesional tissue of mice subjected to cortico-striatal (MCAO) or cortical (PT) stroke. Data are presented as box-and-whisker plots, with the center line indicating the median, + indicating the mean, boxes representing the interquartile range, and whiskers the minimum-to-maximum range. Statistical comparisons were performed using an ordinary two-way ANOVA with brain region and treatment/group as factors, followed by Šídák's multiple-comparisons test. Significant pairwise comparisons are indicated by brackets; exact P values are shown.
